# Rapid polygenic selection generates fine spatial structure among ecological niches in a well-mixed population

**DOI:** 10.1101/2020.03.26.009787

**Authors:** Moritz A. Ehrlich, Dominique N. Wagner, Marjorie F. Oleksiak, Douglas L. Crawford

## Abstract

Evolution by natural selection may be effective enough to allow for recurrent, rapid adaptation to distinct niche environments within a well-mixed population. For this to occur, selection must act on standing genetic variation such that mortality i.e. genetic load, is minimized while polymorphism is maintained. Selection on multiple, redundant loci of small effect provides a potentially inexpensive solution. Yet, demonstrating adaptation via redundant, polygenic selection in the wild remains extremely challenging because low per-locus effect sizes and high genetic redundancy severely reduce statistical power. One approach to facilitate identification of loci underlying polygenic selection is to harness natural replicate populations experiencing similar selection pressures that harbor high within-, yet negligible among-population genetic variation. Such populations can be found among the teleost *Fundulus heteroclitus. F. heteroclitus* inhabits salt marsh estuaries that are characterized by high environmental heterogeneity e.g. tidal ponds, creeks, coastal basins. Here, we sample four of these heterogeneous niches (one coastal basin and three replicate tidal ponds) at two time points from among a single, panmictic *F. heteroclitus* population. We identify 10,861 single nucleotide polymorphisms using a genotyping-by-sequencing approach and quantify temporal allele frequency change within, as well as spatial divergence among subpopulations residing in these niches. We find a significantly elevated number of concordant allele frequency changes among all subpopulations, suggesting ecosystem-wide adaptation to a common selection pressure. Remarkably, we also find an unexpected number of temporal allele frequency changes that generate fine-scale divergence among subpopulations, suggestive of local adaptation to distinct niche environments. Both patterns are characterized by a lack of large-effect loci yet an elevated *total number* of significant loci. Adaptation via redundant, polygenic selection offers a likely explanation for these patterns as well as a potential mechanism for polymorphism maintenance in the *F. heteroclitus* system.

**Author Summary:** Evolution by adaptation to local environmental conditions may occur more rapidly than previously thought. Recent studies show that natural selection is extremely effective when acting on, not one, but multiple genetic variants that are already present in a population. Here, we show that polygenic selection can lead to adaptation within a single generation by studying a wild, well-mixed population of mud minnows inhabiting environmentally distinct locations or niches (i.e. tidal ponds and coastal basins). We monitor allele proportions at over 10,000 genetic variants over time within a single generation and find a significant number to be changing substantially in every niche, suggestive of natural selection. We further demonstrate this genetic change to be non-random, generating mild, yet significant divergence between residents inhabiting distinct niches, indicative of local adaptation. We corroborate a previous study which discovered similar genetic divergence among niches during a different year, suggesting that local adaptation via natural selection occurs every generation. We show polygenic selection on standing genetic variation to be an effective and evolutionarily inexpensive mechanism, allowing organisms to rapidly adapt to their environments even at extremely short time scales. Our study provides valuable insights into the rate of evolution and the ability of organisms to respond to environmental change.

## Introduction

Is evolution by natural selection rampant? Does natural selection lead to adaptation on ecological time scales such that populations adapt within generations or among well-connected demes, allowing them to match their local, heterogeneous and temporally unstable environments [1–6]? The consensus response for several decades used to be negative. Evolutionary adaptation on ecological scales is unlikely due to fundamental limits on the extent of selectively important allelic variation and the rate of adaptive change [7–10]. Yet, an increasing number of natural systems demonstrate adaptation on short temporal and small spatial scales [4]: significant divergence among demes associated with different ecological niches within a single population [11], a few generations of competition altering heritable toepad size and habitat use in anoles [12], repeated anthropogenic pollution resistance [13, 14], seasonal change in heritable thermal tolerance [15, 16], response to local anthropogenic heating [1,5,6], and other ecologically relevant traits [4,17–21]. These observations conflict with the predictions of classic population genetics, and their importance is hotly debated [22–26].

Under a polygenic framework, in which redundancy allows for multiple genetic solutions to effect a phenotypic change, adaptation only requires slight allele frequency changes at a subset of potentially adaptive loci which are already segregating in the population [27–30]. The advantages of redundant, polygenic adaptation are manifold: i) selection on standing variation is highly effective and can occur within a single generation, ii) genetic load is reduced, iii) polymorphism and therefore future adaptive potential is largely maintained, and iv) adaptation is less sensitive to migration since maladaptive alleles are also of small effect [27–33].

However, demonstrating redundant, polygenic adaptation in a natural setting is inherently challenging [30,34,35]. Firstly, phenotypic variance is split among multiple loci, thereby reducing per-locus effect size. This implies allele frequency changes in response to selection will also be minor and difficult to identify and distinguish from neutral drift or demographic effects [22,24,36–39]. Secondly, genetic redundancy implies that the number of allelic variants required to reach a local phenotypic optimum (*n_opt_*) is much lower than the total number of variants affecting a trait (*n_tot_*), i.e. *n_opt_ << n_tot_* [30, 40]. Hence, unique subsets of redundant alleles may equally lead to local adaptation in replicate populations [14,30,34,35,41]. Thirdly, as per-locus, additive effect sizes decrease, gene-by-gene interactions must be increasingly responsible for any phenotypic effect [34,35,41]. Consequently, it is unlikely that natural selection will alter the same loci among replicate populations or in replicate experiments exposed to the same selection pressures, i.e. there is redundancy in the adaptive loci [30]. Distinguishing adaptive allele frequency changes from stochastic processes such as drift will therefore be extremely challenging, especially given the countless theoretical models in which certain parameterizations of demography, mutation rates, and purifying selection are shown to create genomic patterns typically associated with selection on standing genetic variation [22,24,29,37].

We are beginning to explore polygenic selection and redundancy in laboratory settings by employing massively parallel, experimental selection [42]; however studies pursuing redundant, polygenic adaption in the “wild” are rare and mostly human focused [27, 43]. Here, we harness the *Fundulus heteroclitus* model system to demonstrate redundant, polygenic adaptation occurring at extremely small temporal and spatial scales in the wild. *F. heteroclitus*, a small, marine teleost, native to the eastern coast of the USA, primarily inhabits tidal estuaries and demonstrates extremely high site fidelity to a single watershed [44–48]. Within these estuaries exist several, unique niches (or microhabitats), each characterized by distinct biotic and abiotic factors such as temperature, dissolved oxygen, or predator abundance [49–54]. *F. heteroclitus* inhabit the entire estuary and demonstrate some degree of site fidelity to specific niches, thus forming multiple subpopulations within the larger population. Nevertheless, fish from across the estuary reproduce at a common location on a yearly basis [55], essentially homogenizing allele frequencies and maintaining panmixia.

Contrary to the prediction that panmictic breeding every generation would inhibit local adaptation, previous work identified significant genetic divergence among *F. heteroclitus* inhabiting distinct niches less than 100m apart in three replicate estuaries/populations [11]. The authors identified single nucleotide polymorphisms (SNPs) displaying significant spatial differentiation among well-connected niches in each of three isolated estuaries and supported by three different selection tests. While none of the outlier loci were shared among all three replicate populations, many SNPs occur either: i) within the same gene at a different position, ii) in a duplicate gene or paralog, or iii) among genes with similar annotations or narrow, well-defined GO-terms [11]. That is, while no outlier SNPs were shared among the three replicate populations, there were signals of selection in the same gene or genes of similar function, a hallmark of redundancy. The authors concluded that redundant, polygenic selection was surprisingly effective in altering allele frequencies among multiple, distinct SNPs that likely share similar biological functions in response to environmental and ecological differences over very small geographic distances. Yet, it is difficult to believe that the large genetic divergence observed at specific SNPs among well-connected niches is due to local adaptation.

The study presented here builds on prior work and tests for rapid local adaptation within a single, panmictic *F. heteroclitus* population by harnessing both spatial and temporal data. We specifically examine subpopulations residing in distinct niches and quantify temporal allele frequency changes from spring to fall, when natural mortality is highest [56]. We find both the number and magnitude of temporal allele frequency changes to be beyond what would be expected by drift or sampling error alone. Analyses are strengthened further by detecting significant concordance in allele frequency changes among subpopulations. Next, we compare temporal outlier loci to genetic distance between subpopulations and find that allele frequency changes during summer generate significant spatial divergence. While this is indicative of differential selection, we find no loci of large effect and only moderate per-locus signals. Yet, the *total number* of loci that are demonstrating both allele frequency change in time and divergence among niches is significantly elevated. Hence, although the signal is moderate, as would be expected given the limitations of testing this concept, the data presented here support the hypothesis of redundant, polygenic adaptation.

## Results

### Phenotypic disparity yet negligible population structure

Of the 2000 fish tagged in spring 2016, 195 (9.8%) were successfully recaptured in the fall of the same year. Recaptured *F. heteroclitus* demonstrated high site fidelity with 186 (95.4%) individuals recaptured at their respective collection site. Nine migrant fish were excluded from further analysis.

*F. heteroclitus* subpopulations displayed significant length differences in spring (Kruskal-Wallis, *p* << 0.05) (Fig 1A), with basin individuals exhibiting the highest mean total length. The bimodal length distribution of all subpopulations suggests the presence of two cohorts of different ages (S1 Fig). In the basin subpopulation, the proportion of larger and likely older fish is substantially higher than in the ponds. The reason for this asymmetric age structure is unclear but could potentially be due to increased mortality of larger (i.e. older) fish in the ponds. Growth rates, calculated for recaptured individuals only, were also significantly different among locations (Kruskal-Wallis, *p* << 0.05) (Fig 1B). Notably, while basin residents show marginally higher growth rates when grouping individuals from all ponds (Mann-Whitney U, *p* = 0.058), this is driven by significant variation among distinct ponds (Mann-Whitney U, *p* < 0.05). In fact, Pond 1 residents exhibit the same growth rate as basin residents (Mann-Whitney U, *p* = 0.52), suggesting each pond may present unique environmental conditions [57].

**Fig 1.**
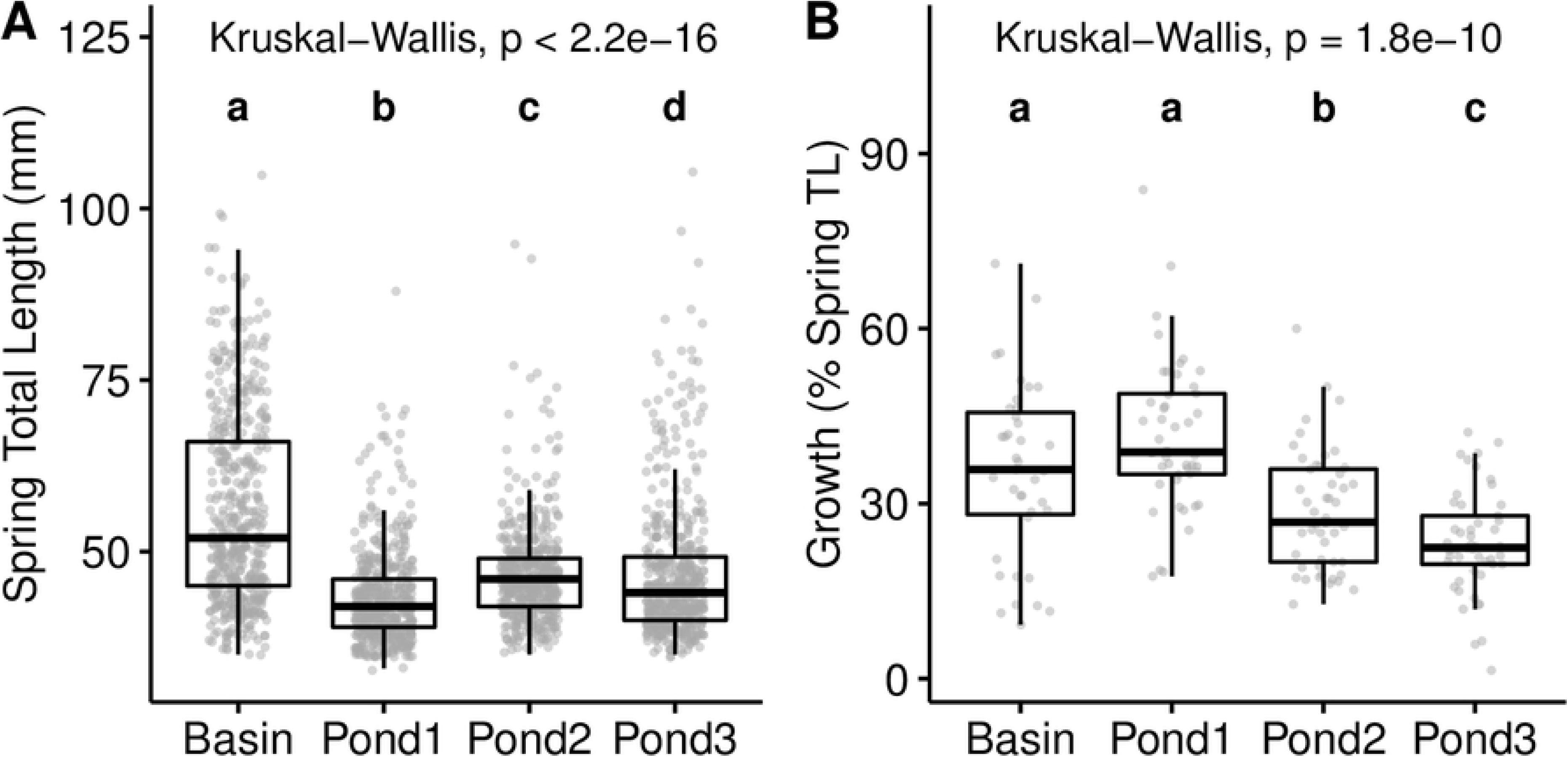
Significant phenotypic differences among subpopulations. Boxplots showing total length of all tagged fish in spring (A) and growth rate of recaptured individuals as a percentage of spring total length (B). Individual data points are shown as gray dots. Global Kruskal-Wallis tests are highly significant for both spring length and growth. Post-hoc, pairwise comparisons are displayed as lowercase letters; subpopulations with the same letter are not significantly different (Wilcoxon, *p* < 0.05).

Despite high site fidelity and marked phenotypic differences among sampling locations, *F. heteroclitus* show negligible population structure based on all 10,861 SNPs, with a global, mean weighted *F_ST_* among all subpopulations and time points of 8.5 x 10^-4^. A PCA biplot using all 10,861 SNPs (S2 Fig) shows negligible structure in both space (i.e. among sampling sites) and time (i.e. between seasons). The absence of spatial structure is expected in a highly connected population with yearly panmictic breeding. Likewise, the lack of a temporal signal between spring and fall collections is indicative of negligible overall change, expected for populations near equilibrium. Consequently, the absence of both spatial and temporal structure suggests that any signal is likely limited to a minor subset of alleles.

### Significant allele frequency changes over time

Resident fish are assumed to have been exposed to their niche-specific environments and associated selection pressures during summer. Any selective death or deterministic emigration will therefore be reflected in significant allele frequency changes relative to the spring collection.

The significance of temporal allele frequency change was quantified by the geometric mean of *p* values generated from three separate significance tests (Barnard’s Test, permutations and simulations) (Fig 2). This approach yielded 611 significant SNPs in the Basin, 664 in Pond 1, 571 in Pond 2 and 625 in Pond 3, each undergoing allele frequency changes that are unlikely due to sampling error, random death or random emigration (geometric mean *p* < 0.05). However, these totals narrowly exceed 543 (5% of 10,861); the expected number of false positives under a uniform *p* value distribution. In fact, only two SNPs within Pond 1 remain significant after multiple test correction (red points, Fig 2). All SNPs that were significant at an FDR of 10% are also significant at the Bonferroni level and are hence displayed as the latter. The absence of major temporal allele frequency changes paired with a moderately elevated *number* of significant SNPs is suggestive of widespread allele frequency changes of small effect, associated with polygenic adaptation [34,35,41].

**Fig 2.**
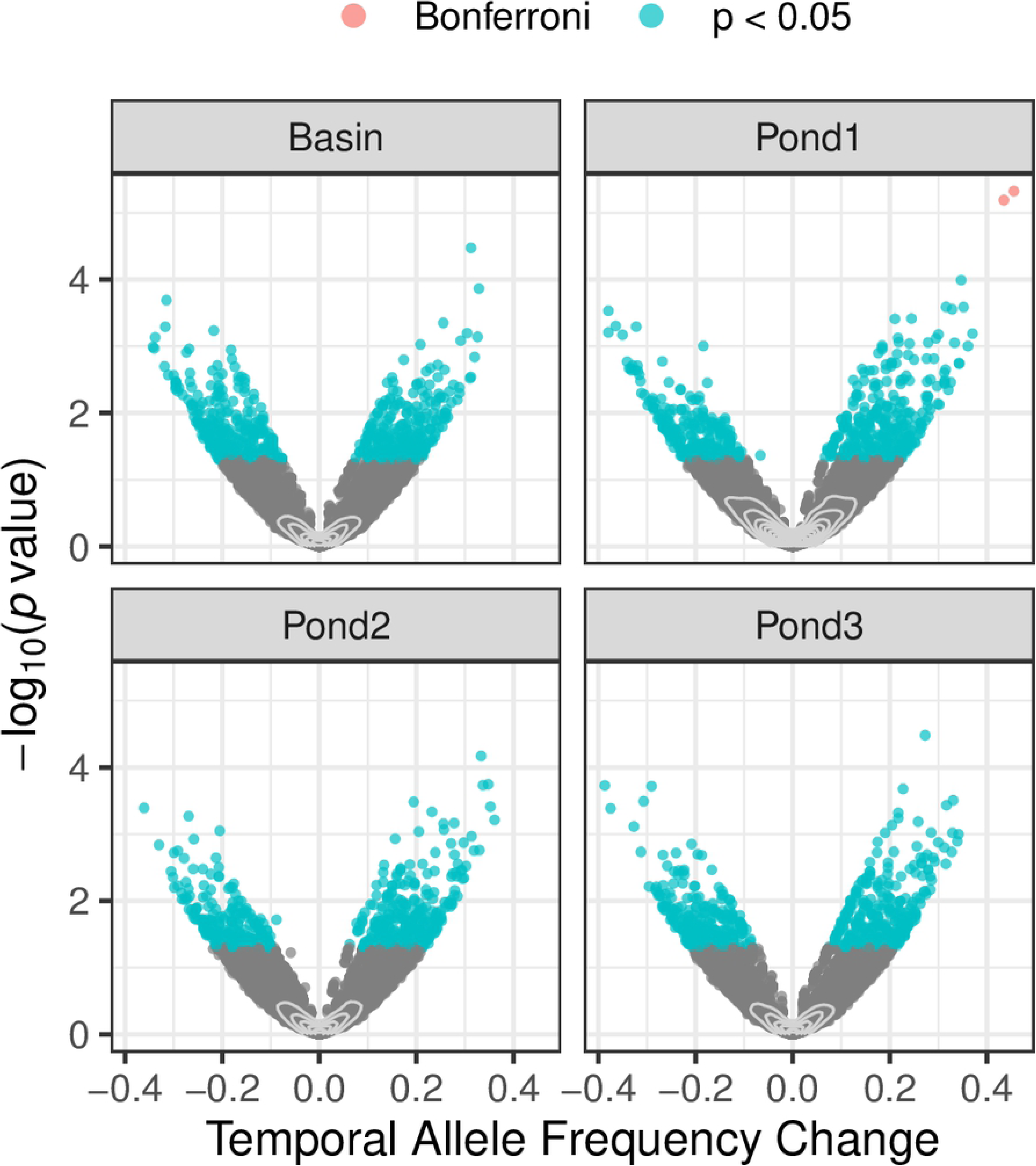
Significant temporal allele frequency change. Temporal change in the reference allele frequency and associated *p* value for all 10,861 SNPs. *p* values shown are the geometric mean of *p* values generated from three separate significance tests (Barnard’s Test, permutation and simulation). SNPs with a significant allele frequency change are shown in color (blue, *p* < 0.05; red, Bonferroni corrected). Grey contour lines show the high density of loci with insignificant allele frequency changes.

To further investigate this idea, the observed *number* of significant SNPs is compared to the expected *number* of false positives assuming a uniform *p* value distribution. Fig 3 shows the observed to expected ratio (O:E) of temporally significant SNPs evaluated across a spectrum of alpha levels to avoid an arbitrary significance threshold. Within every subpopulation, the observed number of significant, temporal allele frequency changes exceeds the expectation for alpha levels between 0.1 and 0.001. Depending on the specific subpopulation, there are approximately 10-50% more SNPs showing significant, temporal allele frequency change than expected due to drift and sampling error. At extreme alpha levels below 10^-3^ (gray shading), both the observed and expected number of significant SNPs are low (<10) causing O:E ratios to become highly discrete and volatile. This complicates interpretation of the data; nonetheless O:E ratios trend above 1. We find both the magnitude and number of significant, temporal allele frequency changes to be independent of niche type, with basin and pond subpopulations showing similar patterns. Yet, while few SNPs remain significant following multiple test correction, it is the total *number* of significant SNPs, each showing minor allele frequency changes, that is unexpectedly high for all subpopulations.

**Fig 3.**
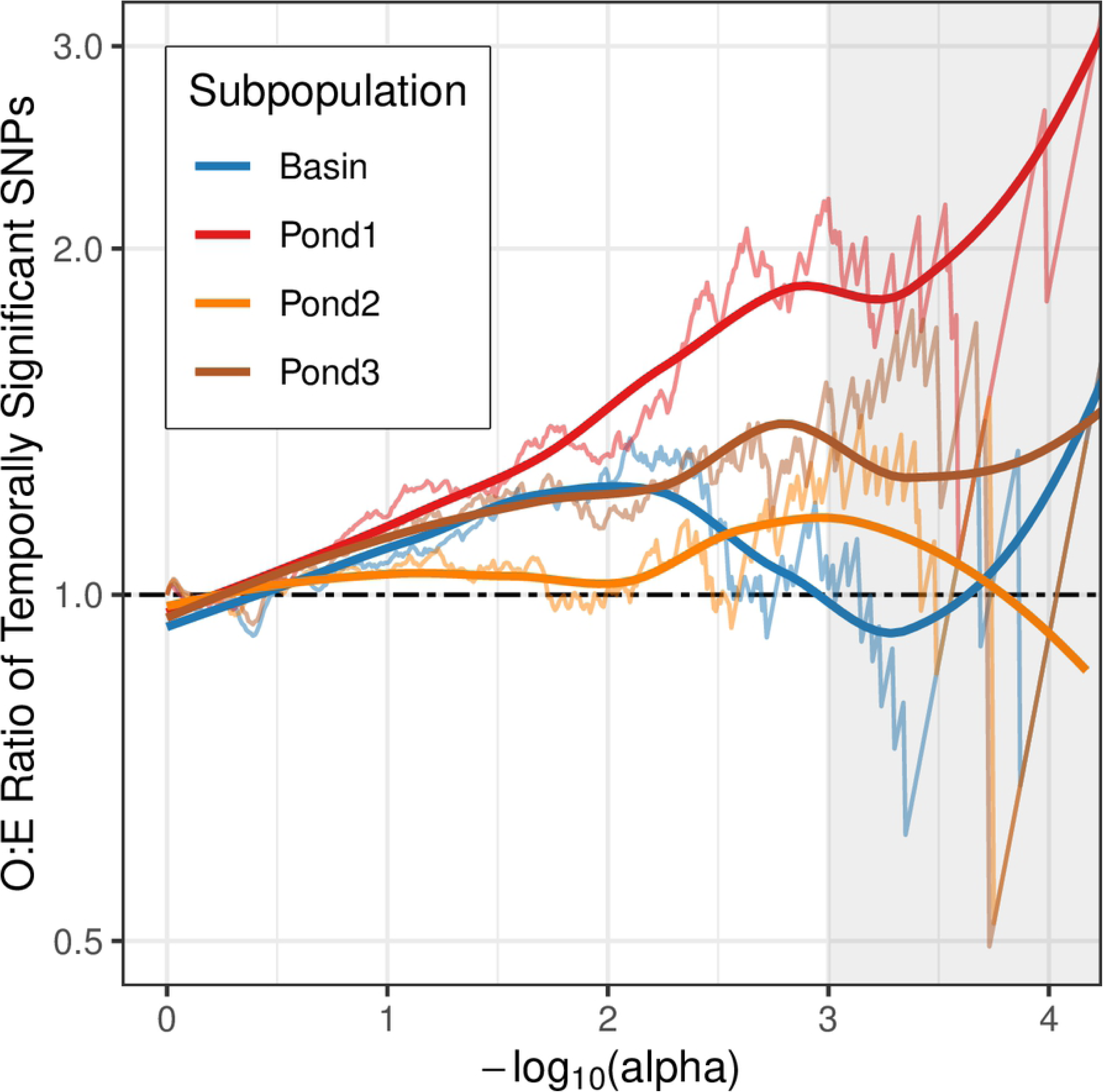
Number of temporally significant SNPs exceeds expectation. Observed to expected ratio of the number of significant temporal SNPs, evaluated across significance thresholds (alpha levels). Thin lines connect O:E ratios, thick lines are splines to aid the eye. The black, dashed line marks the expected ratio of 1 under the null hypothesis. Grey shading marks alpha levels for which the expected number of significant SNPs is 10 or below, giving highly discrete, volatile O:E ratios.

### Concordant allele frequency changes over time

To further test the hypothesis of minor but deterministic allele frequency changes, we assessed the concordance of allele frequency changes in both direction and magnitude, specifically among pond subpopulations, exposed to similar environmental conditions and selection pressures.

Of the 10,861 SNPs tested, we find 3 that show significantly concordant allele frequency changes among pond residents at an FDR of 10% with one showing significance at the Bonferroni level. While this is not overwhelming, pond subpopulations (red line, Fig 4) also display a significantly elevated *number* of concordant allele frequency changes, exceeding both the theoretical expectation based on a uniform *p* value distribution and greatly exceeding 1000 simulations of neutrality for which concordance is spurious by design (gray lines, Fig 4). Simulated data confirms the CMH test is well-behaved and likely conservative, with the smoothed mean of simulations falling well below the null O:E ratio of 1 (black line, Fig 4). In contrast, the O:E ratio for the pond-triplet is 1.6 when evaluated at an alpha level of 10^-2^, corresponding to 60% more SNPs than expected exhibiting concordant allele frequency changes among the three ponds over summer. At alpha levels below 10^-3^ (gray shading) O:E ratios are once again volatile due to the low number of both observed and expected SNPs.

**Fig 4.**
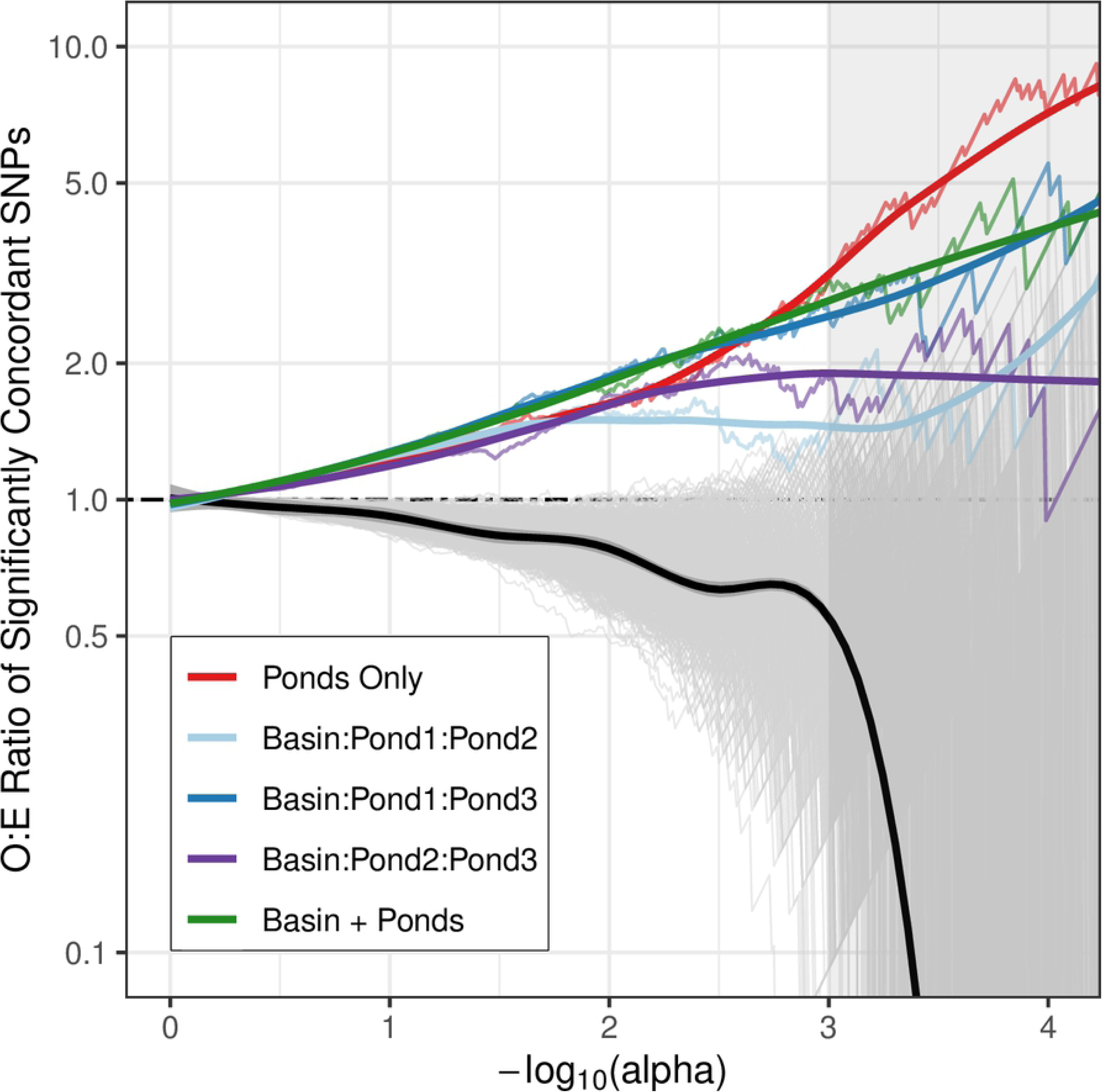
Significant temporal concordance in allele frequency changes among subpopulations. Observed to expected ratio of the number of significantly concordant SNPs among subpopulations (Cochran-Mantel-Haenszel Test). Thin, colored lines connect O:E ratios evaluated across significance thresholds (alpha levels), thick lines are splines to aid the eye. Thin, grey lines represent 1000 simulated subpopulation triplets under the null hypothesis of no temporal change and no spatial differentiation. The solid, black line shows the mean, simulated O:E ratio under the null and its 95% confidence interval. The dashed, black line marks the theoretical expected ratio of 1 under the null hypothesis. Grey shading marks alpha levels for which the expected number of significant SNPs is 10 or below, resulting in highly discrete, volatile O:E ratios.

While the unexpectedly large number of concordant SNPs among pond subpopulations is suggestive of niche-specific allele frequency change as a result of parallel selection, other triplets which include the basin subpopulation show similarly elevated O:E ratios (blue shaded lines, Fig 4). In fact, the number of concordant SNPs among all subpopulations (green line, Fig 4) is up to 5-fold higher than expected and largely exceeds 1000 neutral simulations, indicative of mutual, ecosystem-wide allele frequency shifts. Of these, 2 SNPs remain significant after multiple test correction at an FDR of 10% with one showing significance at the Bonferroni level. Allele frequency changes at these loci seem to be unrelated to niche type and possibly due to a common selection pressure experienced by all subpopulations.

### Fine spatial structure among subpopulations

While there is negligible spatial structure when utilizing all 10,861 SNPs (S2 Fig), pairwise comparisons among subpopulations in fall yield individual loci with exceptional *F_ST_* values (Fig 5). In fact, SNP-specific *F_ST_* values often exceed 0.2, ranging as high as 0.43. Nevertheless, only two pairwise comparisons display SNPs that remain significant after multiple test correction; Basin:Pond 2 (2 SNPs) and Pond1:Pond2 (1 SNP). Again, all SNPs that are significant at an FDR of 10% are also significant at the Bonferroni level and are hence displayed as the latter. Furthermore, pond:pond pairwise comparisons yield *F_ST_* outliers with similar magnitudes to basin:pond comparisons, contrary to the prior expectation of genetic divergence among distinct niches [11].

**Fig 5.**
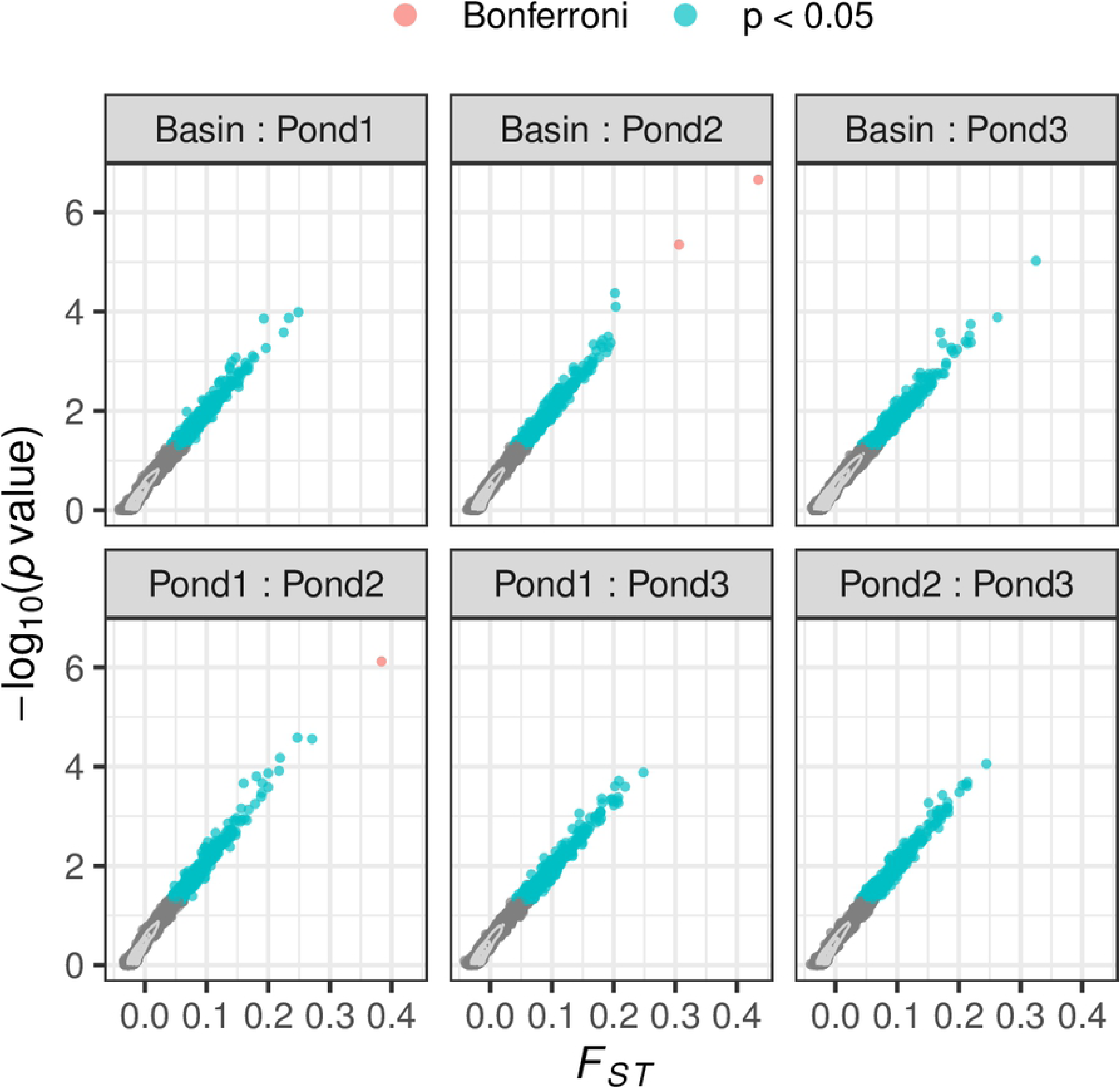
Significant spatial outlier SNPs in fall. *F_ST_* and respective *p* value for all spatial pairwise comparisons evaluated at all 10,861 SNPs in fall. Basin:Pond pairwise comparisons are displayed in the top row, Pond:Pond comparisons in the bottom row. *p* values shown are the geometric mean of *p* values generated from three separate significance tests (Barnard’s Test, permutation and simulation). SNPs with a significant *F_ST_* are shown in color (blue, *p* < 0.05; red, Bonferroni corrected). Grey contour lines mark the high density of SNPs with insignificant *F_ST_*.

The results presented here partially corroborate the findings of Wagner *et al.* [11] who found significant spatial outlier SNPs among niches in this and other salt marsh estuaries. However, the authors did not compare within niche type, making it difficult to attribute spatial divergence to an niche effect as opposed to divergence among subpopulations distributed in space. Additionally, of the 4,741 SNPs assayed by Wagner *et al.* [11], only 115 are successfully genotyped here. The low overlap can be mostly attributed to the filtering required to optimize data completeness as well as the thinning approach employed by both studies to minimize inflating outlier numbers due to linkage disequilibrium. Nevertheless, of the 63 spatial outlier SNPs identified by Wagner *et al.* [11] in 2013, 4 are present in the current data set of which one shows significant spatial divergence (basin:pond 1, *p* < 0.05).

Perhaps more important than the significance of individual, spatial outlier SNPs is the proportion of SNPs exhibiting significant spatial differentiation as compared to the expectation under complete panmixia. Fig 6 shows the ratio of the observed *versus* expected number of spatially significant SNPs, evaluated along a continuum of alpha levels. For every pairwise comparison the O:E ratio of spatially significant SNPs falls above 1 for moderate alpha levels above 10^-3^. Hence, a significant number of SNPs display spatial differentiation beyond what is expected due to neutral processes, indicative of fine, spatial structure among subpopulations. Contrary to prior expectations [11] but consistent with temporal data, pond:pond pairwise comparisons yield similar, significantly elevated numbers of spatially diverged SNPs (mean of 635 significant SNPs per pairwise comparison, *p* < 0.05) as opposed to basin:pond comparisons (mean of 638 spatially significant SNPs per pairwise comparison, *p* < 0.05).

**Fig 6.**
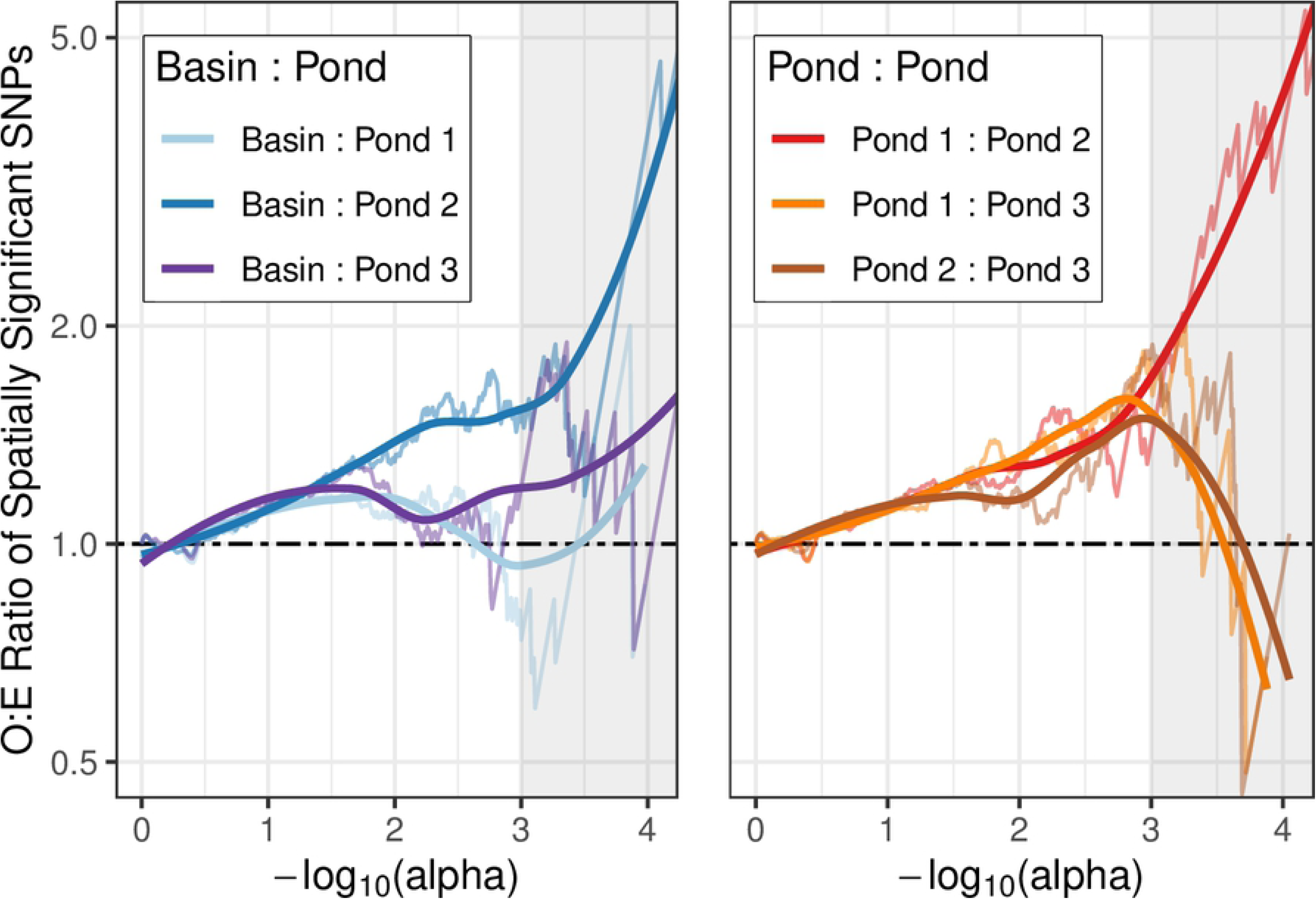
Number of spatial outlier SNPs exceeds expectation. Observed:Expected ratio of the number of spatially significant SNPs, evaluated across a spectrum of alpha levels for each pairwise comparison among subpopulations in fall. Basin:Pond pairwise comparisons are displayed on the left, Pond:Pond comparisons on the right. Thin lines connect computed O:E ratios, thick lines are splines to aid the eye. Black dotted lines mark the expected ratio of 1 under the null hypothesis of zero spatial differentiation. Grey shading marks alpha levels for which the expected number of significant SNPs is 10 or below, giving highly discrete, volatile O:E ratios.

However, both the absence of loci with large *F_ST_* as well as the lack of covariance between genetic divergence and environment is expected under polygenic selection with redundancy [30,34,35,41]. Pond subpopulations may be phenotypically differentiating from the basin each using a unique set of redundantly adaptive alleles. Such adaptation with redundancy would result in parallelism at the phenotype level yet present as mild divergence at multiple, distinct loci. Alternatively, cryptic environmental variation may be the cause of distinct phenotype optima leading to divergence among pond subpopulations [57].

### Recent allele frequency changes generate fine spatial structure

Both the number of loci exhibiting a significant temporal signal, as well as those displaying significant spatial divergence, exceed the neutral expectation in this system. In order to elucidate the spatiotemporal relationship of these non-neutral patterns, we examined the intersection of temporally and spatially significant SNPs and found a significant enrichment of SNPs that are both changing in time and differentiated in space (Chi-squared, *p* << 0.01). This holds true upon evaluating all temporally significant SNPs, regardless of subpopulation, and all spatially significant SNPs, regardless of pairwise comparison. Significant enrichment also occurs within any given pairwise comparison i.e. when only temporally and spatially significant SNPs pertaining to the subpopulations in a specific pair are taken into account (Chi-squared, *p* << 0.01 for all pairwise comparisons). This significant enrichment suggests that temporal allele frequency changes within subpopulations generate fine spatial structure among subpopulations.

Fig 7 displays how magnitude and direction of temporal allele frequency changes affect spatial differentiation between subpopulation pairs. To simplify visualization, data from all six pairwise comparisons were collapsed into a single graphic. The x- and y-axis display temporal allele frequency change for any two arbitrary subpopulations while the coloration indicates the spatial differentiation among these subpopulations in fall (see S3 Fig for specific pairwise comparisons). At SNPs where subpopulations have undergone large, opposing (antagonistic) allele frequency changes (top-left and lower-right quadrant, Fig 7), spatial divergence is highest. SNPs exhibiting no or parallel (concordant) allele frequency changes between subpopulations (top-right and lower-left quadrant, Fig 7) display low levels of differentiation, mostly because prior (minor) allele frequency differences remain unchanged. As expected, the majority of SNPs are centered on the origin, displaying neither significant temporal change, nor spatial differentiation among subpopulations (gray contour lines, Fig 7). The circular symmetry indicates allele frequency changes are mostly uncorrelated among subpopulation pairs. The distribution of spatially differentiated SNPs (yellow shading, Fig 7) suggests that antagonistic allele frequency changes during the summer months are primarily responsible for fine spatial structure observed in fall.

**Fig 7.**
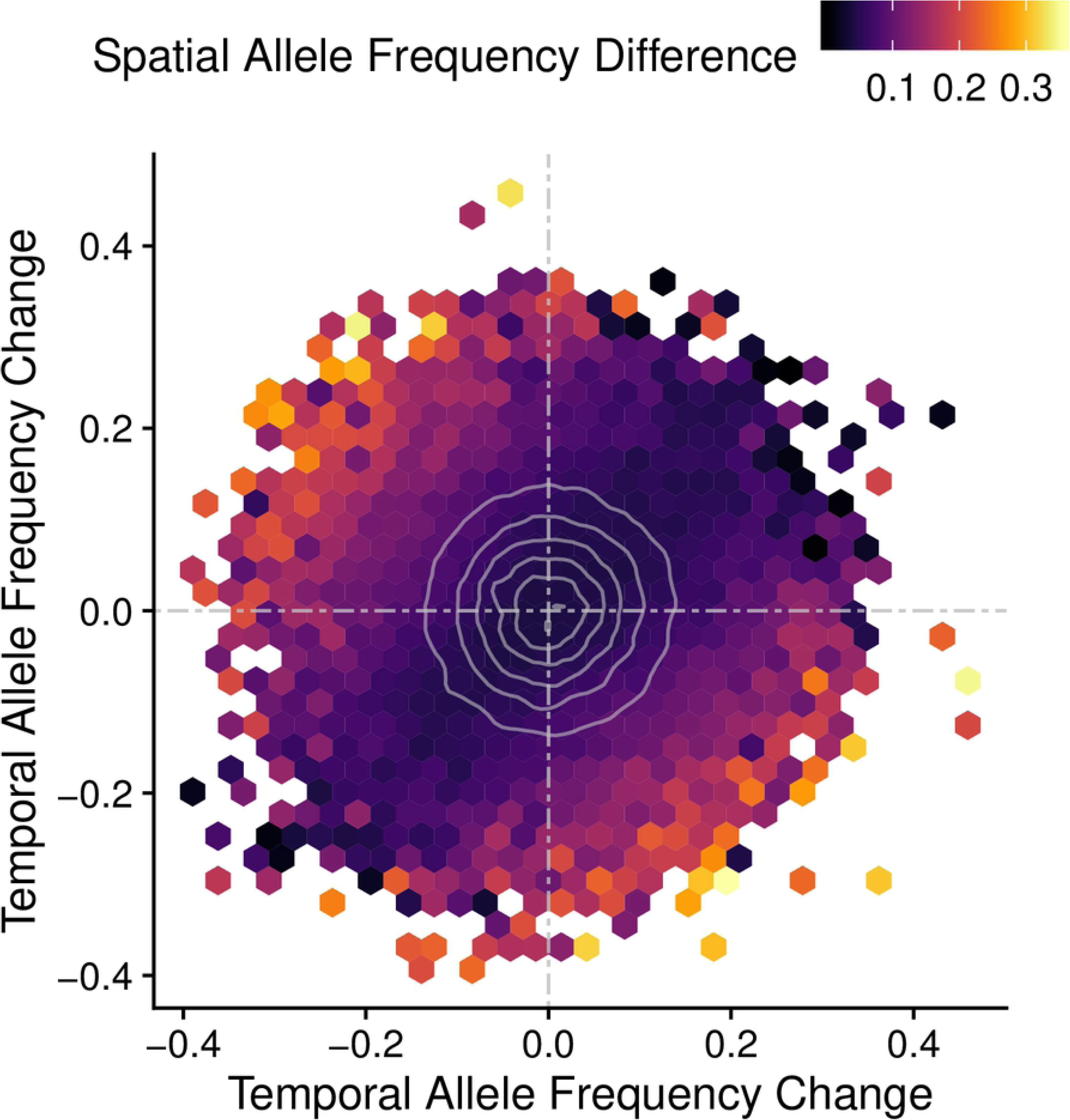
Antagonistic allele frequency changes generate spatial allele frequency differences among subpopulations. Heatmap showing the relationship between temporal allele frequency changes in two subpopulations (x- and y-axis) and the mean spatial allele frequency difference between these subpopulations in fall (coloration). All pairwise comparisons are overlaid with subpopulations arbitrarily assigned to either the x-or y-axis. Dotted, gray lines mark the origin; solid, gray contour lines the density distribution of SNPs.

To test this concept, Fig 8 shows posterior (fall) spatial allele frequency differences as a function of prior (spring) spatial allele frequency differences (A) and contrasts this to posterior (fall) spatial allele frequency differences as a function of relative temporal allele frequency change (B) (see methods for definition of relative temporal allele frequency change). Panel A shows no correlation between prior and posterior spatial allele frequency differences. While significantly positive, both effect size and variance explained by prior spatial structure are negligible (*p* << 0.01, adj. *R^2^* < 10^-3^). On the contrary, panel B shows a significant, positive correlation between relative allele frequency change in time and posterior (fall) spatial allele frequency differences (*p* << 0.01, adj. *R^2^* = 0.42).

**Fig 8.**
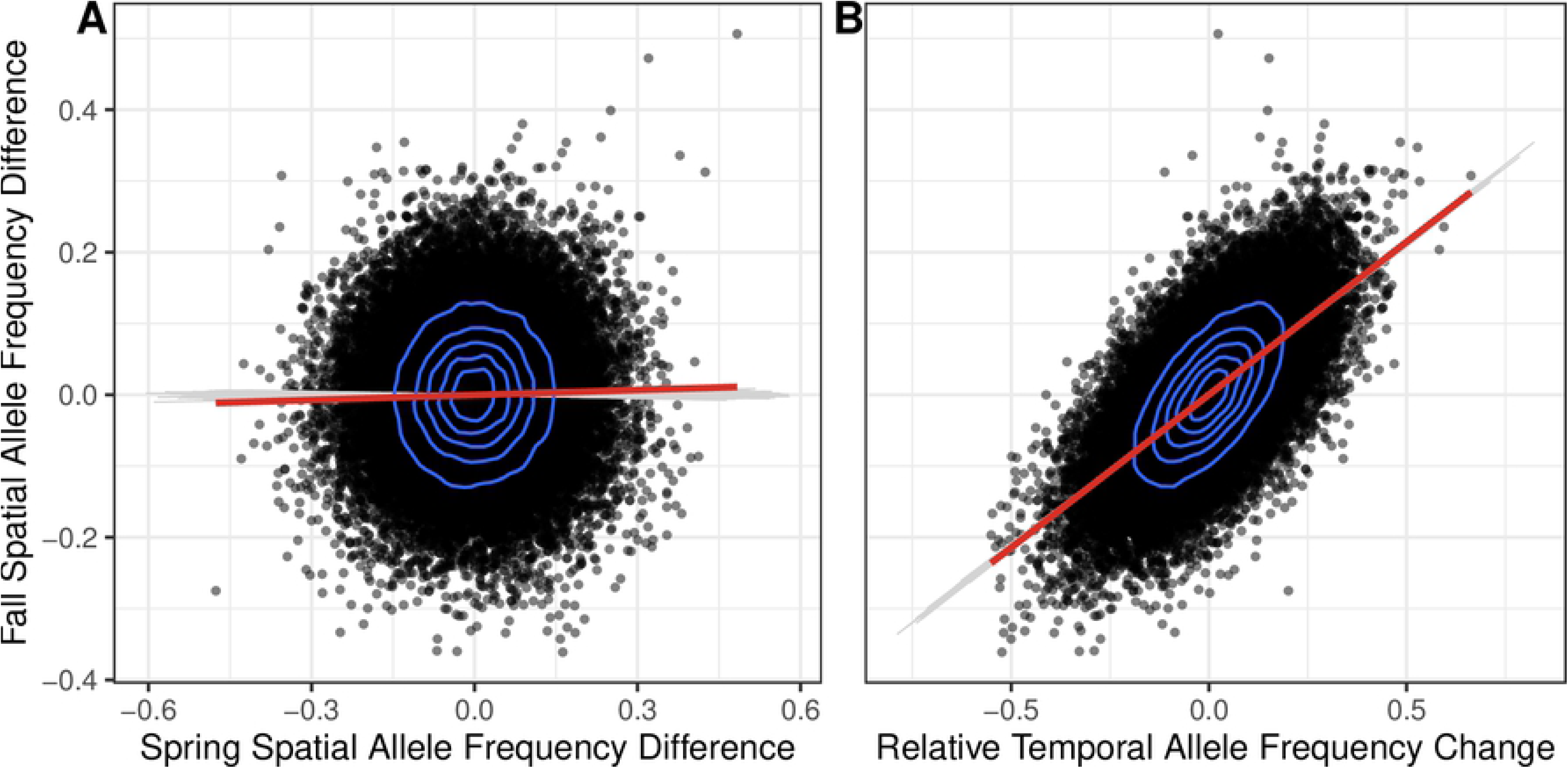
Recent allele frequency changes, not prior structure, explain fine spatial structure in fall. A) Scatterplot showing no correlation between spatial allele frequency differences in spring versus fall (red line, *p* << 0.01, adj. *R^2^*<10^-3^). B) Scatterplot showing a significant, positive correlation between the relative allele frequency change of a subpopulation pair and the respective spatial allele frequency difference observed in fall. The red line shows a linear regression of the empirical data (*p* <<0.01, adj. *R^2^*= 0.42). Data from all pairwise comparisons is overlaid. Blue contour lines mark the high density of SNPs near the origin. Gray lines are linear fits of 1000 simulated data sets of spatially homogeneous and temporally invariant subpopulations.

These patterns highlight two major insights. Firstly, large, relative allele frequency changes, i.e. the joint allele frequency change of a subpopulation pair due to the shift of one subpopulation relative to the other, generate large, spatial allele frequency differences (Fig 8B). Essentially, antagonistic allele frequency changes (i.e. large, relative changes) produce spatially divergent SNPs, whereas parallel changes (i.e. small relative changes) rarely result in significant spatial differences. Secondly, and more importantly, temporal allele frequency changes are more predictive of posterior (fall) allele frequency differences (i.e. fine spatial structure) than prior (spring) structure. In other words, recent, temporal allele frequency changes generate fine spatial differentiation among subpopulations.

## Discussion

### Non-neutral patterns in time and space

High site fidelity at small spatial scales [44–48] combined with a large tag-and-recapture effort allowed for *in situ* monitoring of selective processes in this *F. heteroclitus* population. The disparate temperature and dissolved oxygen regimes [49, 50] experienced by tidal pond and coastal basin residents are plausible drivers of selection given their direct effect on fitness related life-history traits in ectotherms [50,58–64].

Minimal spatial structure (S2 Fig) throughout the estuary provides a homogeneous genetic baseline upon which selection may act in any single environmental niche. In fact, recently acquired GBS data of larval fish, caught throughout the estuary 2-4 weeks post-spawning, does not show any population structure, nor increased kinship among larvae caught at the same location (unpublished). This is highly suggestive of panmictic breeding and random dispersal, yet we find significant morphological differences among fish inhabiting distinct locations within the estuary (Fig 1).

We find an elevated number of SNPs that undergo significant temporal allele frequency changes from spring to fall, i.e. changes that are unlikely due to random death, sampling effects, or other neutral processes alone (Fig 3). In addition, we find an unexpectedly high proportion of allele frequency changes to be concordant among subpopulations in both magnitude and direction (Fig 4). Spatial data corroborates previous findings by Wagner *et al.* [11], showing an elevated number of significantly differentiated SNPs among interbreeding, resident subpopulations within the same estuary in fall (Fig 6). Finally, we show that loci undergoing significant temporal changes also exhibit high spatial differentiation, suggesting that spatial structure in fall is primarily determined by allele frequency changes taking place during the summer months, not by prior spatial structure in spring (Fig 7).

### Temporal change and spatial divergence are confounded

Rapid adaptation to a heterogeneous environment via selective death from a common baseline offers an explanation for the covariance between temporal allele frequency change and spatial allele frequency differences. Selective death presents as unexpectedly large allele frequency changes at effector SNPs within each subpopulation, essentially reconfiguring spatial structure according to these shifts. The genetic landscape is now a direct result of recent allele frequency changes at the ecological scale. SNPs that discriminate among subpopulations appear to have recently undergone allele frequency changes, resulting in a correlation among spatially and temporally significant SNPs. Such an evolutionary scenario appears to provide a parsimonious explanation for the surprising, fine-scale structure observed among salt-marsh subpopulations here and demonstrated previously in three different *F. heteroclitus* estuaries [11].

While it is tempting to attribute the enrichment of SNPs that are both temporally and spatially significant to rapid adaptation, inconsistent spatial structure through time and high explanatory power of recent, temporal changes can also be a consequence of neutral processes. In fact, temporal allele frequency change and spatial allele frequency difference within the same population are inherently confounded.

To demonstrate this concept, we performed 1000 simulations of four spatially homogeneous and temporally invariant subpopulations, each based on the empirical allele frequency distribution. Only neutral processes, such as random death and sampling effects were permitted to generate allele frequency changes, and hence, apparent spatial structure. The correlation between these neutral, temporal allele frequency changes and resulting spatial allele frequency differences can be seen in Fig 8B. For clarity, individual SNPs have been omitted and only the linear regression line has been plotted for each simulation (gray lines, Fig 8). Simulation regression lines fall within a tight range and show a near identical spatiotemporal relationship as the empirical data. In fact, both the empirical correlation coefficient and adjusted *R^2^*-value fall within one standard deviation of the mean simulated values.

This result demonstrates the inherent relationship between recent, temporal allele frequency change and current spatial structure. By constructing an equation defining posterior spatial allele frequency difference as a function of prior spatial difference, and relative temporal allele frequency change between two populations, the inherent relationship becomes apparent:

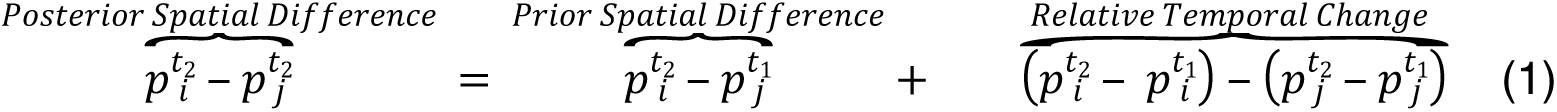

Where the super- and subscript denote time point and population respectively. In the light of equation (1) one can see that the residuals of the linear regression in Fig 8A are equivalent to the relative, temporal allele frequency changes. Similarly, the residuals in Fig 8B are equivalent to the prior, spatial allele frequency differences. The spatiotemporal correlation shown in Fig 8B is therefore a direct consequence of this mathematical relationship and will present itself in any system where temporal allele frequency changes are large and dominate prior, spatial allele frequency differences. Given its negligible spatial structure, this is precisely the case in the *F. heteroclitus* system.

We acknowledge that the inherent relationship between temporal change and spatial structure cannot be used as an argument for selection since it holds true regardless of whether neutral or deterministic processes generate allele frequency change. However, we have provided ample evidence to demonstrate that the allele frequency changes and associated spatial structure observed here at multiple loci cannot be explained by neutral processes alone. Specifically, we have shown that an unexpectedly large number of SNPs display significant allele frequency changes in every subpopulation. We have also demonstrated there to be significant concordance in allele frequency changes among subpopulations. Finally, we corroborate a previous study by presenting a significant number of SNPs that are substantially diverged among subpopulations in fall. We therefore propose local adaptation via polygenic selection as the most parsimonious explanation, leading to deterministic allele frequency changes that have generated ecologically meaningful spatial sub-structure in the *F. heteroclitus* system.

### Parallel selection across the ecosystem

An unexpectedly high number of concordant allele frequency changes among all subpopulations (basin and all three ponds) is a clear signal of non-neutral processes and suggestive of a global selection pressure, affecting the entire estuary. Given the prolonged presence of *F. heteroclitus* in New Jersey salt marshes (∼15,000 years) [65, 66], it is unlikely for these concordant allele frequency changes to be the result of continued, long-term adaptation to a distant trait optimum. Large effective population sizes [67, 68] should allow globally advantageous alleles to become fixed over such a time span, especially when selection is strong enough to produce allele frequency changes on the order of those shown here. Instead, we suggest adaptation to a recent change in phenotypic optimum as a potential driver of ecosystem-wide, concordant allele frequency changes.

Temporal heterogeneity in biotic or abiotic factors such as predator abundance [21, 69] or temperature [70] could lead to a common, cyclic selection pressure in all *F. heteroclitus* subpopulations. If the period of these environmental fluctuations is on the order of 3 years or less (approximate *F. heteroclitus* life-span in the wild [71]), then overlapping generations could lead to global, concordant allele frequency changes via the storage effect [72–75]. Briefly, the storage effect describes an evolutionary mechanism in which part of the population is protected from selection during a specific life stage, thereby “storing” non-beneficial alleles until environmental conditions revert to the point where these become beneficial again [75]. A multitude of organisms experience such age-specific selection, with early-life stages often being the most vulnerable [76, 77]. Fish like *F. heteroclitus* are no exception, displaying highest mortality (and likely selection) at the larval stage [56,71,78]. Adult fish may be protected from selection, despite “storing” deleterious alleles that do not match current conditions. Yearly reproduction by the adults propagates “stored” alleles until the environment reverts to conditions under which these alleles confer increased fitness, allowing them to rise in frequency once again. Additionally, the storage effect is further stabilized by high plasticity in the adults [79]. If adults can readily tolerate periods unfavorable to their genotype through plasticity, they are more likely to propagate their alleles during future, favorable conditions. *F. heteroclitus* is known to exhibit exceptional plasticity and is often used as a model organism for studying individual, phenotypic variance [1,80–82].

Hence, we suggest that a genetic storage effect that maintains polymorphism in the population and allows for a rapid adaptive response to an environmental change via selection from standing genetic variation is a plausible explanation for the significantly concordant, ecosystem-wide allele frequency changes observed here.

### Genetic divergence among subpopulations

The hypothesized niche effect, in the form of differential allele frequency changes in pond versus basin subpopulations leading to divergence between these two niches, did not present itself. Nevertheless, we observe significant phenotypic differences and an unexpected number of significantly differentiated loci among resident subpopulations, regardless of niche type. This surprising result, previously demonstrated by Wagner *et al.* [11], suggests the presence of another ecologically relevant effect that is not necessarily rooted in pond/basin environments.

Environmental heterogeneity is unlikely to present as discrete partitioning into binary niche types, but rather as a continuum resulting from the combined effect of multiple, possibly obscure, environmental factors. Stuart *et al.* [57] have shown that cryptic environmental differences among stickleback lake and stream habitats can explain this apparent lack of parallelism among populations pairs. Similarly, potential cryptic environmental variables among ponds may be of similar, or higher, importance as the documented temperature and dissolved oxygen differences. For example, Hunter *et al.* [50, 51] demonstrated that tidal pond flooding frequency is significantly, positively correlated with female *F. heteroclitus* gonadosomatic index. The authors suggest that increased nutrient availability, introduced by frequent tidal flooding, may allow for higher reproductive allocation, a key life-history and fitness related trait. Consequently, quasi-isolated, resident subpopulations throughout the saltmarsh may be exposed to an unknown, heterogeneous fitness landscape with distinct selection pressures resulting in unique pheno- and genotypic responses.

We propose two independent and inclusive mechanisms that may be responsible for the observed moderate yet significant divergence among resident subpopulations in a heterogeneous environment: antagonistic selection and matching habitat choice.

#### Antagonistic selection in a heterogeneous environment

Levene [83] first discussed the consequences of a spatially heterogeneous landscape on the distribution and maintenance of polymorphism in a population. In his model, a finite number of demes inhabit distinct ecological niches. Selection takes place within each niche, leading to a local increase of alleles conferring higher fitness in the respective environment. Next, individuals from every niche panmictically breed, with offspring randomly dispersing into niches and selection commencing again. If two alleles at the same locus each confer increased fitness to one niche, then both alleles will be maintained in proportion to the relative contributions of each niche to the global population. At the phenotype level this can be imagined in terms of a trade-off for a specific trait among niches, in which each allele produces a phenotype that is advantageous in one and deleterious in another environment. In this manner a patchy environment i) may lead to significant allele frequency changes within niches as selection proceeds and ii) could maintain polymorphism in the global population [84].

Estuaries inhabited by *F. heteroclitus* consist of multiple niches with contrasting environmental parameters. In addition, cryptic differences among seemingly similar niches further exacerbate environmental heterogeneity [49–51,64]. Given this patchy landscape and annual panmictic breeding, it is possible that both the significant temporal allele frequency changes as well as spatial divergence observed in the *F. heteroclitus* system could be explained by Levene’s model. Niches across the estuary have distinct phenotype optima with locally advantageous alleles increasing in frequency due to selective death. Survivors then reproduce panmictically in the upper intertidal and larvae disperse at random throughout the estuary, reinitiating the process. Alleles that are universally advantageous are likely fixed over time while antagonistic alleles that exhibit a tradeoff among niches are maintained in the global population.

While Levene’s model seems to provide a plausible explanation for the observed patterns, it has several underlying assumptions. Specifically, it requires i) antagonistic effects to be equal in magnitude to prevent fixation of the fitter allele [85, 86], ii) environment-dependent reversal of dominance [87, 88] and iii) minimal migration among niches to avoid swamping with maladapted alleles [89].

While these limitations are restrictive, the *F. heteroclitus* system described here mostly fulfils and/or alleviates these assumptions. Firstly, high phenotypic plasticity in *F. heteroclitus* [1,80–82] may dampen the necessity of balanced antagonism [79]. Secondly, relative fitness within niches outweighs imbalanced antagonism in global fitness if niches are resource-limited [83, 90]. This seems to be the case in the small bodies of water inhabited by *F. heteroclitus* [50,51,56,64]. Hence, the relative contribution of an allele to the next generation is proportional to the carrying capacity of its matching niche, not relative global fitness [83, 90]. Thirdly, reversal of dominance is conceivable and has been successfully demonstrated in the lab [91, 92]. If antagonistic alleles are deleterious recessive [93, 94], or pleiotropic [94], which is likely the case for redundant alleles affecting complex traits, dominance reversal can be readily achieved. Finally, while *F. heteroclitus* does not fulfil the stabilizing requirement of negligible migration among habitat patches [44, 51], intermediate levels of migration can potentially exacerbate local adaptation. If migration is deterministic, i.e. individuals actively seek niches that best match their genotype, the proportion of beneficial alleles within niches becomes inflated further stabilizing Levene’s model [83]. Matching habitat choice is a possibility in the *F. heteroclitus* system and in fact complimentary to antagonistic selection.

#### Matching habitat choice

The majority of *F. heteroclitus* exhibit extremely limited dispersal (>60% stay within 20 meters of tagging location), yet a significant proportion readily travels between niches during spring high tides when the estuary floods [44,45,47,48]. For example, during early summer, pond emigration rates can reach up to 30% per month compared to mortality rates of 20% [51]. Similar migration rates have also been reported in the basin, suggesting niches are in fact highly connected [44]. The tag-and-recapture approach employed in this study cannot discriminate between mortality and emigration. Although only a small proportion of individuals migrated among sampling sites (4.6%), we cannot exclude the possibility of emigration into unsampled locations. Consequently, the significant temporal allele frequency changes and spatial differences observed here are likely to be the result of deterministic mortality and/or deterministic emigration.

Non-random, genotype-dependent gene flow, referred to here as matching habitat choice, is an often overlooked alternative to local adaptation when presented with genotype-environment covariance [95–98]. Significant, spatial genetic divergence may in fact be the result of individuals “self-sorting” into niches, by actively sensing their surroundings and seeking environments in which their fitness is maximized [98]. In contrast to the classic interpretation of migration which homogenizes genotypes among demes, matching habitat choice promotes heterogeneity. Such directed gene flow can tilt migration-selection balance in favor of selection, leading to local adaptation in the broad sense even when migration is unfeasibly high for classic, narrow sense adaptation (i.e. local adaptation via natural selection) [97]. Furthermore, matching habitat choice significantly reduces genetic load in outbred populations and can account for polymorphism maintenance since alleles are essentially redistributed in space, not removed through selective death [96].

While several studies, both in the lab and field, have demonstrated matching habitat choice to be the prevalent mode of local adaptation in the broad sense [99–103], in most cases it is a costlier adaptive strategy compared to phenotypic plasticity and/or selection [95, 96]. In fact, matching habitat choice is only favorable in in an actively dispersing species [104] that breeds panmictically, with offspring distributing randomly throughout a habitat that is highly heterogeneous both in space and time yet offers minimal barriers to dispersal [96, 98]. *F. heteroclitus* inhabiting New Jersey salt marshes evidently meet these conditions, making matching habitat choice a plausible, alternative explanation for the observed, significant phenotypic differences, elevated number of temporal allele frequency changes, and spatial divergence among subpopulations.

### Redundant polygenic selection

Both the total number of significant temporal allele frequency changes and spatial allele frequency differences are beyond the neutral expectation, yet only few loci remain significant following multiple test correction. We acknowledge that large-effect loci may have been missed since the GBS approach only queried approximately 0.3% of the 1 Gb *F. heteroclitus* genome. In addition, linkage disequilibrium is negligible, extending to approximately 1 Kb (S4 Fig), making it unlikely for genotyped SNPs to be linked to rare, large-effect loci. Nevertheless, finding any significant signal under these circumstances is highly improbable unless such signals are pervasive.

An unexpectedly high number of loci undergoing non-neutral changes, yet each with minor effect size is consistent within a framework of polygenic selection in which trait variance is split among multiple loci [34,35,41]. Especially complex, fitness-related life-history traits have been shown to be highly polygenic [58,105–107]. It is therefore plausible that the significant, minor allele frequency changes observed at multiple loci are the result of soft selective sweeps on complex traits [30, 108]. Likewise, spatial divergence presents itself as subtle allele frequency differences among subpopulations at multiple loci [30,34,35,41].

While we detect significantly differentiated loci among distinct niche types (basin and ponds), we also observe a similar number and magnitude of divergent SNPs among ponds. This may be the consequence of local adaptation to uncategorized, environmental variation among apparently analogous niches [57], e.g. flooding frequency [50]. Alternatively, it may reflect redundancy in the adaptive alleles required to achieve increased fitness i.e. redundancy in genotype-to-phenotype mapping [30,34,35,41]. Replicate populations experiencing similar selection pressures and adapting towards a common phenotypic optimum may therefore not share the same adaptive alleles [30,40,42]. Further, genetic redundancy offers a potential explanation for the maintenance of polymorphism in the *F. heteroclitus* system, which would allow for soft sweeps to occur on standing variation every generation [30].

Although we detect unexpected divergence among subpopulations inhabiting similar niches, we also find significant concordance in allele frequency change among subpopulations. This is likely to occur if genetic redundancy is limited, increasing the probability of the same loci presenting in replicate subpopulations. A second possibility is that the effect size distribution of loci underlying a polygenic trait is not uniform. In other words, certain loci may have a larger effect on phenotype, experience stronger selection and be detected more easily in replicate populations [109, 110](but see [111]).

### Conclusion and future directions

We have discovered significant, concordant allele frequency changes among independent subpopulations of a well-mixed, larger *F. heteroclitus* population that are likely the result of ecosystem-wide adaptation to a common phenotype optimum. At the same time, we find temporal allele frequency changes that generate fine, yet significant, divergence among subpopulations, suggesting local adaptation to distinct niche environments. Antagonistic selection and matching habitat choice are potential, complimentary mechanisms that can explain these patterns. Finally, while local adaptation via redundant, polygenic selection is not unambiguously proven here, it offers a conceivable explanation for the lack of large-effect loci yet elevated *number* of significant loci as well as a potential mechanism for polymorphism maintenance in the *F. heteroclitus* system.

Nevertheless, further work is necessary to validate this interpretation. Firstly, comprehensive phenotyping of individuals following summer conditions will be required to confirm selection is altering trait means among niche residents. Given the stark differences both in temperature and dissolved oxygen levels between pond and basin habitats [49], traits known to be impacted by these abiotic factors are likely to be the most promising choices [50,58–64]. Next, testing for differential survivorship in reciprocal transplants of basin and pond residents could confirm trait divergence is due to prior selection on niche conditions, not acclimation. Caging may be used to eliminate matching habitat choice as a potential mechanism.

Comprehensive whole genome sequencing is required to confirm redundant, polygenic architecture in *F. heteroclitus*. Reduced-representation sequencing and low levels of linkage disequilibrium severely limit our ability to detect potential large-effect loci. Only by sequencing the entire genome can the effect-size distribution and genetic architecture be inferred. A repeat of this experimental design may then allow for comparisons between years i.e. are allele frequency changes and the resulting spatial divergence consistent among years? Finally, by combining phenotypic and genetic data, associations can be drawn between alleles displaying significant frequency changes in a given niche and the respective phenotype under selection. Such association mapping, potentially using polygenic scores, may then shed light on the process of rapid niche adaptation in a well-mixed population.

Detecting subtle signatures of redundant, polygenic selection on complex traits remains elusive. We acknowledge the limitations of the data presented here but encourage further work addressing rapid, polygenic adaptation, not only in a laboratory setting, but within wild populations.

## Methods

### Tagging and sample collection

Initial tagging and tissue collection took place in late spring 2016 (22 May – 5 June) at the Rutgers University Marine Field Station (RUMFS), NJ. Over a 10-day period, 2200 *F. heteroclitus* were caught using minnow wire traps at four sampling sites (550 fish per site) throughout a single saltmarsh estuary (Fig 9). The collection sites included a coastal basin and three permanent, intertidal ponds, all part of the same watershed and interconnected during spring tides occurring approximately 5-15 times per month [50, 51].

**Fig 9.**
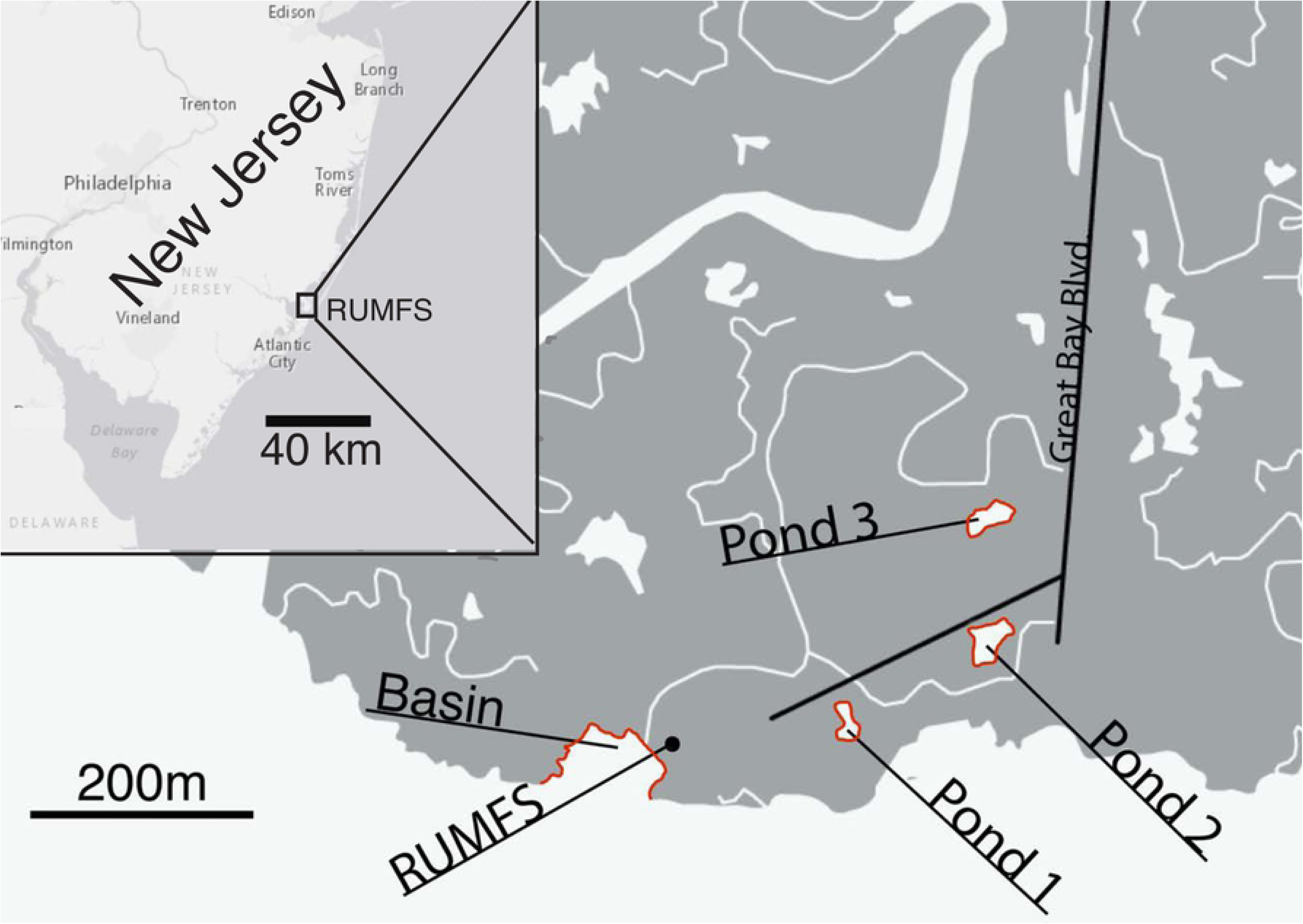
Location of the four sampling sites within the watershed around the Rutgers University Marine Field Station (RUMFS). Insert shows the location of the RUMFS on the coast of New Jersey, USA.

500 fish from each sampling site were weighed, measured (total length), sexed and uniquely tagged using sequential coded wire tags (Northwest Marine Technology Inc.). Caudal fin clips were taken from the remaining 50 fish and stored in guanidinium hydrochloride (GuHCl) buffer solution [112]. After tagging/clipping, fish were released at their respective capture location. Tagged fish were recaptured in early fall 2016 (30 August – 10 September) by trapping at the same 4 collection sites. Trapping efforts were continued until 50 tagged fish had been recaptured at each location (only 45 were recaptured in the basin). Recaptured, tagged fish were weighed, measured (total length) and sexed. Caudal fin clips were taken from all 200 recaptured fish and stored in GuHCl buffer solution. Coded wire tags were dissected from each individual, then read and cross-referenced with spring tagging data. Only residents, i.e. fish that were tagged and recaptured at the same location, were included in further analysis. This sampling scheme allowed for spatial comparisons among sampling sites as well as assessment of temporal change from spring to fall. While summer residency does not implicate genetic, ecological or reproductive substructure, for the purpose of this analysis, resident individuals from a single collection site are henceforth collectively referred to as a subpopulation. Differences in length and growth rate among subpopulations were tested using Kruskal-Wallis ANOVAs followed by post-hoc, Mann-Whitney U tests in *R* v3.6.1 [113]. Weight data was excluded due to the high abundance of gravid females during spring collection that may have confounded results.

### DNA isolation and library preparation

Genomic DNA was isolated from 30 individuals from each resident subpopulation and time point (spring and fall) using a custom SPRI magnetic bead protocol, yielding a total of 240 isolates. Genotyping-by-sequencing (GBS) libraries were prepared using a modified protocol after Elshire et al. (2011). In short, high-molecular-weight genomic DNA was aliquoted and digested using the AseI restriction enzyme. Digests from each sample were uniquely barcoded, pooled and size selected to yield insert sizes between 350-550 bp. Pooled libraries were PCR amplified using custom primers that extend into the insert by 1 base (cytosine). This approach systematically reduces the number of sequenced tags, ensuring sufficient sequencing depth.

### Sequencing, SNP calling and filtering

Pooled libraries were sequenced on one lane of the Illumina HiSeq 4000 in 2×150 bp paired-end mode yielding approximately 467 million paired-end reads (>140 Gb). Single nucleotide polymorphism (SNP) calling was performed using the GBeaSy analysis pipeline [115] with the following filter settings: minimum read length of 30bp after barcode and adapter trim, minimum phred-scaled variant quality of 30 and minimum read depth of 5 at the sample level. This yielded a total of 3,775,496 SNPs that were further filtered using VCFtools 0.1.13 [116]. Specifically, only biallelic SNPs without significant heterozygote excess (tested at p-corrected <0.01 [117]) were included. Furthermore, SNPs were filtered to a maximum of 10% missing data and a minimum minor allele frequency (MAF) of 5%. Filtering was applied at the individual level to only include samples with more than 60% completeness i.e. <40% missing genotypes per sample. Finally, the variant set was thinned to a minimum of 300bp between SNPs on the same scaffold. This guaranteed a single SNP per read, minimizing linkage disequilibrium among SNPs. The final, filtered set contained 193 individuals genotyped at 10,861 SNPs on which all further analysis is based. Sample sizes by location and season remained relatively balanced following filtering (spring:fall = 86:107; basin:pond1:pond2:pond3 = 51:46:49:47). Here, ‘allele frequency’ refers to the frequency of the *F. heteroclitus* reference genome allele. Since only biallelic SNPs were used in the analysis, the alternate allele frequency is implied.

### Temporal allele frequency change and spatial differentiation

Global and pairwise *F_ST_* statistics were calculated using VCFtools 0.1.13 [116]. Principal component analysis (PCA) was performed in *R* v3.6.1 [113] using the *SNPRelate* package v1.18.1 [118].

*P* values relating to spatial (i.e. among sites) and temporal (i.e. between seasons) comparisons were attained via three separate, yet dependent tests. These included 1) a permutation approach, 2) comparison to a spatially or temporally homogeneous simulated population and 3) a Barnard’s exact test. Permutations were performed using a custom, parallelized bash script (GitHub address pending). *P* values were generated by comparing empirical *F_ST_* values to permuted values conditioned on heterozygosity as in *FDIST2* [119]. Samples from all locations and both time points were included in the analysis in order to increase permutational space and decrease the likelihood of generating sets that closely match empirical data.

Simulations were performed in *R* v3.6.1 [113] where sets of subpopulations were generated *in silico* under the null hypothesis of zero temporal change. Specifically, spring and fall empirical data were considered two samples of a temporally invariant subpopulation. For every SNP, the weighted mean allele frequency of the spring and fall samples was used to estimate the allele frequency under the null hypothesis. This null allele frequency was then used to generate a simulated subpopulation in Hardy-Weinberg equilibrium for every SNP. Simulated subpopulations were then randomly sampled *n* times without replacement, where *n* is the empirical sample size at each SNP. SNP-specific Hardy-Weinberg subpopulations were sampled twice, representing the empirical spring and fall sample. Random sampling of the simulated subpopulations is representative of random death in the wild as well as experimental sampling effects, both in the field and during sequencing. Finally, the simulated spring allele frequency was subtracted from the simulated fall allele frequency to obtain temporal allele frequency change. The above sampling procedure was repeated 10,000 times, generating SNP-specific distributions of apparent allele frequency change under the null hypothesis of zero temporal change. Empirical allele frequency changes were then compared to the simulated distributions and *p* values generated according to rank. Separate temporal simulations were produced for every subpopulation in order to account for possible spatial heterogeneity. Simulated subpopulations sizes were 1300 and 400 for the basin and ponds respectively. These estimates are in agreement with the observed subpopulation sizes in the wild based on exhaustive sampling at each collection site.

Simulations testing spatial structure were conducted in a similar fashion. Here SNP-specific global populations were generated under the null hypothesis of spatial homogeneity. Subpopulations were considered samples of a single, large, panmictic, global population and their weighted mean used to estimate the global neutral or null allele frequency. Empirical allele frequency differences among sites were compared to the distribution of apparent allele frequency differences under the null and *p* values generated as above. Spatial simulations were only conducted on fall data in order to remain agnostic to possible temporal change.

Barnard’s exact test was performed on contingency tables of allele counts comparing either temporal changes (within each subpopulation) or pairwise spatial differences (fall only). Barnard’s test is statistically similar to Fisher’s Exact test and, whilst computationally more costly, better suited to genetic data since it does not condition on margin totals. Tests were performed using the *Exact* package [120] in *R* v3.6.1.

To facilitate further analysis, *p* values from permutations, simulations and Barnard’s tests were combined by taking their geometric mean. The geometric mean is a conservative aggregate metric, appropriate for combining correlated *p* values from dependent tests [121, 122]. This resulted in a single *p* value per SNP and subpopulation, quantifying the significance of temporal change, as well as a single *p* value per SNP and pairwise comparison, quantifying the significance of spatial differentiation in fall. Multiple test correction was performed using the *p.adjust* function in *R* v3.6.1 [113] by applying both the false discovery rate [123] and Bonferroni methods.

### Temporal concordance

A Cochran-Mantel-Haenszel (CMH) test [124, 125] was applied to temporal data in order to determine whether significant allele frequency changes were concordant among niches, specifically ponds, and hence likely due to selection. The CMH test assesses the degree of concordance with respect to both magnitude and direction of allele frequency change. Primarily, concordance among the three replicate pond subpopulations was tested whilst other “triplet” comparisons, comprised of two ponds and the basin, served as inherent controls. A CMH test evaluating concordance among all four subpopulations was also conducted. In addition, CMH tests were performed on 1000 simulations of temporally invariant subpopulations (see above for details) to assess the degree of spurious concordance. Tests were performed on allele counts using the *mantel.haenszel* function from the *base* package in *R* v3.6.1 [113].

### Spatiotemporal correlation

In order to elucidate the relationship between temporal change and spatial structure, a linear model was constructed in which posterior (fall) spatial allele frequency difference was regressed onto prior (spring) allele frequency differences. Relative temporal allele frequency change, Δ*p_ij_*, between a pair of populations is defined as:

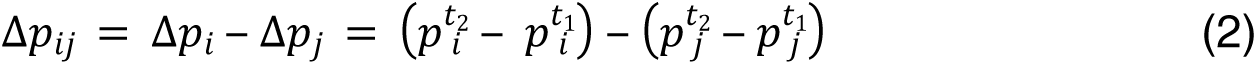

Where the super- and subscript denote time point and population respectively. This metric quantifies the degree to which the allele frequencies of two populations converge or diverge over time. The two regression analyses allowed for comparing the relative explanatory power of prior spatial structure as opposed to recent temporal change in determining posterior spatial structure. Regression analyses were performed for every pairwise-comparison, then aggregated. In order to evaluate the significance of spatiotemporal correlation in the context of evolutionary adaptation, 1000 simulations of temporally invariant and spatially homogeneous subpopulations were performed (see above for details). As with the empirical data, linear models were fit to both temporal and prior spatial data for each simulation. Regression analyses were performed using the *lm* function in *R* v3.6.1 [113].

### Ethics Statement

Fieldwork was completed within publicly available lands, and no permission was required for access. *F. heteroclitus* does not have endangered or protected status, and small marine minnows do not require collection permits for non-commercial purposes. Adult *F. heteroclitus* were captured in minnow traps with minimal stress and removed in less than 1 hour. Tag-and-recapture and non-surgical tissue sampling protocols were in compliance with and approved by the University of Miami Institutional Animal Care and Use Committee (IACUC, protocols 16-124 and 19-119).

## Acknowledgements

We would like to thank Dr. Kenneth Able, Roland Hagan and all technicians at the Rutgers University Marine Field Station for their discussions regarding mummichog ecology and practical support in the field respectively. Further thanks to Keenan Berry, Katy Erceg, Sathvik Palakurty and Rebecca Pelofsky for field work assistance. Many thanks to Amanda DeLiberto and Melissa Drown for countless discussions regarding the analysis of spatial and temporal genetic data.

---

**S4 Fig. Linkage Disequilibrium decays rapidly.** Weighted, mean correlation coefficient (*r^2^*) among genotypes as a function of physical distance. The blue line marks the smoothed mean correlation coefficient and the gray shading its 95% confidence interval.

**S1 Fig. Age-structure explains size differences among basin and ponds.** Size distributions (total length) by subpopulation of fish tagged in spring 2016 (top) and of recaptured, resident individuals in fall 2016 (bottom).

**S2 Fig. No apparent population structure in space or time.** Principal component analysis of all subpopulations and both time points using all 10,861 SNPs. Biplot of first and second principal component with variance components given in parentheses. Colors represent subpopulations, symbols the sampling time point. 95% confidence ellipses are drawn around the sample means. Dotted lines represent spring, solid lines fall samples. While two outlier individuals segregate from the main cluster along the first two principal components, this pattern does not indicate structure *per se* but rather the inexistence of a principle component that can partition a meaningful amount of variance. That is, the variance due to these outliers is in fact negligible compared to the total variance in the data set.

**S3 Fig. Antagonistic allele frequency changes generate spatial allele frequency differences among subpopulations.** Heatmaps showing the relationship between temporal allele frequency changes in two subpopulations (x- and y-axis) and the mean spatial allele frequency difference between these subpopulations in fall (coloration). The first subpopulation in the facet label was assigned to the x-, the second subpopulation to the y-axis. Dotted, gray lines mark the origin; solid, gray contour lines the density distribution of SNPs.

